# Convergence of MCR-8.2 and chromosome-mediated resistance to colistin and tigecycline in an NDM-5-producing ST656 *Klebsiella pneumoniae* from a lung transplant patient

**DOI:** 10.1101/2021.07.21.453302

**Authors:** Jiankang Zhao, Ziyao Li, Yulin Zhang, Xinmeng Liu, Zhujia Xiong, Yanyan Fan, Xiaohui Zou, Binghuai Lu, Bin Cao

## Abstract

For infection caused by NDM-5-producing *Klebsiella pneumoniae*, tigecycline and colistin are the last treatment options. In this study, we characterized the first NDM-5 and MCR-8.2 co-harboring *K. pneumoniae* clinical isolate, combining with chromosomal gene-mediated resistance to colistin and tigecycline. The *K. pneumoniae* KP32558 was isolated from the bronchoalveolar lavage fluid from a lung transplant male patient. Whole genome sequencing was carried out using Illumina HiSeq sequencing platform as well as nanopore sequencing method. The *K. pneumoniae* KP32558 was identified as a pan-drug resistant bacteria, belonged to ST656, and harbored plasmid-encoded *bla*_NDM-5_ and *mcr-8.2* genes. The *bla*_NDM-5_ gene was located on an IncX3 type plasmid, which was successfully transferred to Escherichia coli strain J53 without visible fitness cost. The *mcr-8.2* gene was located on a conjugative plasmid pKP32558-2-mcr8. It had two replicons, IncFII(K) and IncQ1, harbored previously by another two *mcr-8.2*-carrying plasmids pMCR8_020135 and pMCR8_095845. These three plasmids were clustered into the same clade and derived from K. pneumoniae isolates of the same clonal complex, indicating that pKP32558-2-mcr8 may come from pMCR8_020135 and pMCR8_095845 related ancestor. The MIC of KP32558 for colistin was 256 mg/L, the 6 amino acid substitutions in the two-component system may involve in the high-level colistin resistance. The truncation in *acrR* gene, related to tigecycline resistance, was also identified. *K. pneumoniae* has evolved a variety of complex resistance mechanisms to the last-resort antimicrobials, close surveillance is urgently needed to monitor the prevalence of this clone.

---

The emergence and worldwide dissemination of carbapenem-resistant *Klebsiella pneumoniae* (CRKP) have posed a great threat to public health (1). According to the results of China antimicrobial surveillance network (CHINET) in 2018, the resistance rate of *K. pneumoniae* to imipenem and meropenem has exceeded 25% (2). The main resistance mechanism for CRKP is the production of carbapenemases, in which New Delhi metallo-β-lactamase (NDM) is capable to hydrolyze all β-lactams except monobactam. The *bla*_NDM-5_ gene was first identified from an *Escherichia coli* isolate in 2011 from a patient with a hospitalization history in India (3). Afterward, NDM-5-producing *K. pneumoniae* isolates have been detected and caused sporadic outbreaks worldwide (4, 5). Compared to NDM-1, NDM-5 has two amino acid substitutions at Val88Leu and Met154Leu, showing increased resistance to carbapenems and broad-spectrum cephalosporins when expressed under its native promoter (3).

For infection caused by NDM-5-producing CRKP, tigecycline and colistin are the last treatment options (6, 7). In recent years, colistin resistance in *Enterobacterales* isolates has been found more frequently. Several molecular mechanisms have been associated with colistin resistance, such as chromosomal mutations in *PmrA/PmrB, PhoP/PhoQ*, and *CrrA/CrrB* two-component systems (TCS), and inactivation of the regulator *mgrB* gene (8, 9). Besides, since the first plasmid-encoded colistin resistance gene, *mcr-1*, was identified from *E. coli* in 2015 (10), nine variants of *mcr-1* (*mcr-2* up to *mcr-10*) have been identified in *Enterobacterales* (11). The horizontal transferability of plasmid facilitates the rapid dissemination of colistin resistance. Furthermore, increasing consumption of tigecycline has been concurrent with increasing reports of tigecycline resistance. The main mechanism is the overexpression of resistance-nodulation-cell division (RND)-type efflux pumps (AcrAB and OqxAB) and the mutations in efflux pump regulator genes (*ramA*, *soxR*, *marR*, and *acrR*) (12–14).

The *K. pneumoniae* strains, presenting extensively resistance to carbapenem, tigecycline and colistin simultaneously, would present a great threat to public health. In this study, we isolated a *K. pneumoniae* strain from the bronchoalveolar lavage fluid (BALF) specimen of a lung transplant patient. Antimicrobial susceptibility testing indicated that this strain was a pan-drug resistant *K. pneumoniae*. Revealing the mechanism of drug resistance will play a positive role in preventing the spread of this strain.

## MATERIALS AND METHODS

### *K. pneumoniae* isolates and the reference plasmids

Two *K. pneumoniae* isolates (KP31166 and KP32558) were recovered from the BALF specimens of a lung transplant patient 1 and 4 months after surgery, as shown in Figure 1. For comparative genomic analysis, ten *mcr-8.2*-harboring plasmids were collected from the NCBI genome database, including plasmids pMCR8_020135 (CP037964), pMCR8_095845 (CP031883), pVNCKp115 (LC549807), pZZW20-88K (CP058962), p2019036D-mcr8-345kb (CP047337), pD120-1_83kb (CP034679), pVNCKp83 (LC549808), pSCKLB555-4 (CP043936), p2018C01-046-1_MCR8 (CP044369) and p18-29mcr-8.2 (MK262711).

**Figure 1.**
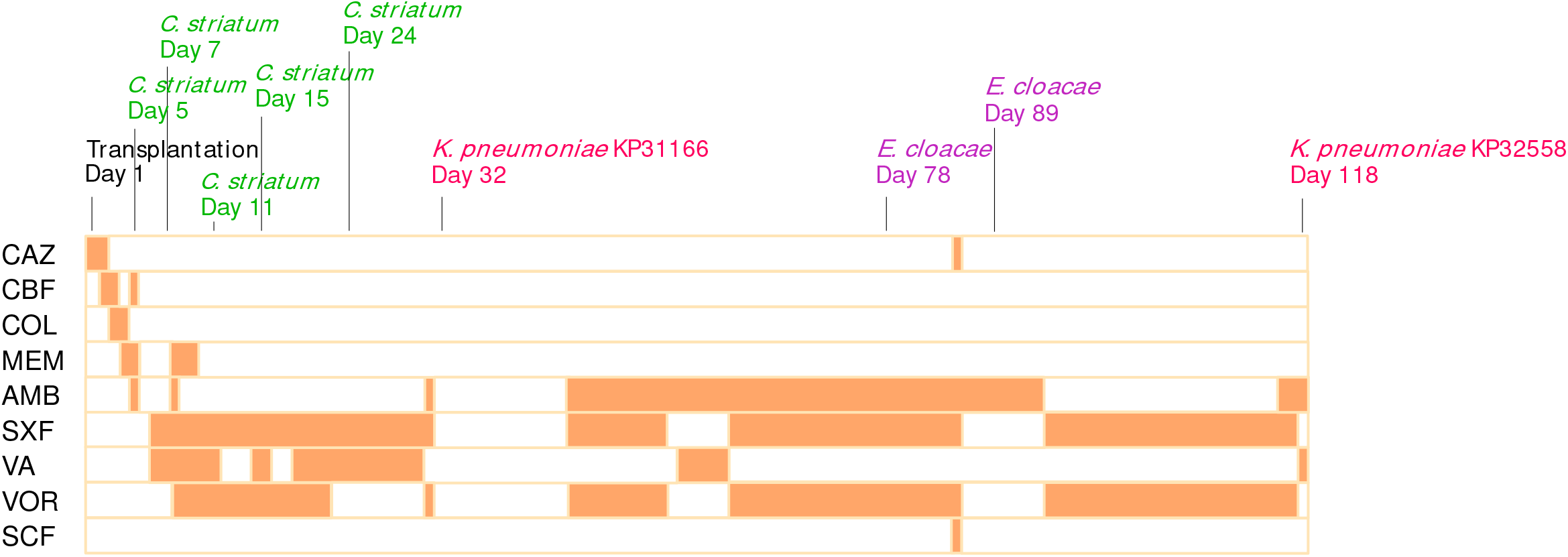
Clinical data of the lung transplant recipient. CAZ, ceftazidine; CBF, caspofungin; COL, colistin; MEM, meropenem; AMB, amphotericin B; SXF, sulfamethoxazole/trimethoprim; VA, vancomycin; VOR, voriconazole; SCF, cefoperazone/sulbactam.

### Antimicrobial susceptibility testing

In vitro susceptibility tests of amikacin, tobramycin, minocycline, doxycycline, ceftazidime, cefepime, piperacillin/tazobactam, cefoperazone/sulbactam, aztreonam, imipenem, meropenem, levofloxacin, ciprofloxacin, and sulfamethoxazole/trimethoprim were performed using Vitek-2 system in N335 susceptibility cards. The minimum inhibitory concentrations (MICs) of tigecycline and colistin were determined using the microdilution broth method, with its reagents provided by bio-KONT, Ltd. China. The production of carbapenemase was determined using the modified carbapenem inactivation method (mCIM) and EDTA-modified carbapenem inactivation method (eCIM) as recommended by the Clinical Laboratory Standards Institute (CLSI) (15). The breakpoint of tigecycline and colistin were defined by the European Committee on Antimicrobial Susceptibility Testing (EUCAST, version 11.0, http://www.eucast.org/). In particular, the minimum inhibitory concentration (MIC) > 0.5 mg/L and > 2 mg/L will be defined as resistant to tigecycline and colistin, respectively.

### Hypermucoviscous phenotype determination

The string test was used to identify the hypermucoviscous phenotype of *K. pneumoniae* strains. Strains exhibiting a viscous string > 5 mm were considered hypermucoviscous (16).

### Whole-genome sequencing (WGS) and annotation

For *K. pneumoniae* isolate KP32558, complete genome sequencing was carried out using Illumina HiSeq 2500 sequencing platform as well as nanopore sequencing method on MinION flow cells. Raw reads were filtered to remove low-quality sequences and adaptors using skewer (17) and PoreChop (https://github.com/rrwick/Porechop), respectively. *De novo* assembly was conducted using SPAdes Genome Assembler v3.13.1 (18) and Unicycler (19). Gene prediction for 11 genomes, including one from this study and 10 from NCBI genome database, was performed using Prokka 1.12 (20). Genomic islands were predicted using IslandViewer 4 (21). Insertion sequences were identified using ISfinder database (22). The antimicrobial resistance genes, multilocus sequence types (MLST) and plasmid replicon were analyzed via the CGE server (https://cge.cbs.dtu.dk/services/). Virulence genes were identified using the BIGSdb *Klebsiella* genome database (http://bigsdb.Pasteur.fr/klebsiella/klebsiella.html). The heatmap was generated using an in-house R script.

### Phylogenetic analysis

A total of 11 *mcr-8.2*-carrying plasmids were included and aligned to reference plasmid pKP32558-2-mcr8 (GenBank accession no. CP076032, this study) using Snippy (https://github.com/tseemann/snippy). A single nucleotide polymorphism (SNP) maximum likelihood (ML) tree was constructed by using FastTree 2 (23) with general time reversible (GTR) model of nucleotide substitution and a gamma distribution of rate heterogeneity. Phylogenetic clusters were identified using hierarchical Bayesian analysis of population structure, in which the first level of clustering was used to define clusters (24). The phylogenetic tree was annotated by using FigTree v1.4.3 (http://tree.bio.ed.ac.uk/software/figtree/).

### Plasmid conjugation assays

The plasmid conjugation experiment was performed for *K. pneumoniae* strain KP32558, and azide-resistant *E. coli* J53 was used as the recipient strain. Transconjugants were selected on LB plates containing azide (100 mg/L) and meropenem/colistin (2 mg/L). Antimicrobial susceptibility testing, PCR amplification and S1-PFGE were performed to confirm the successful transfer. The transfer frequency was calculated as the number of transconjugants per recipient.

### Growth kinetics

Growth curves for the recipients *E. coli* J53 and transconjugants were measured as follows: LB broths containing 1 × 10^6^ CFU/mL bacteria were incubated overnight at 37 °C with consecutive shacking. Meanwhile, the same broth without bacteria was set as the growth control. OD600 measurement was taken at 0, 2, 4, 6, 8, 10, and 12 h to construct a growth curve. All experiments were performed in duplicate.

### Ethics approval

Permission for using the information in the medical records of the patient and the *K. pneumoniae* isolates for research purposes was granted by the Ethics Committee of the China-Japan Friendship Hospital (2019-164-K113).

## RESULTS

### Clinical data of the patient

The patient was a 66-year-old male, who was given colistin to prevent infection 2-3 days after lung transplantation, and the BALF culture was repeatedly negative. The *K. pneumoniae* isolates KP31166 and KP32558 were detected in his BALF specimens on day 32 and 4 months later, respectively. The patient produced a large amount of sputum and had necrotic tissue in the bronchial opening, but the pathogenic detection of necrotic tissue was negative. In addition, the patient had no fever, and blood routine and C-reactive protein tests were normal when *K. pneumoniae* was detected. Considering the condition of the patient and antimicrobial susceptibility results of the *K. pneumoniae*, no specific anti-infective treatment was given (Figure 1). However, *K. pneumoniae* was not isolated from BALF specimens since then, and the patient recovered well.

### General phenotypic and genotypic characteristics

According to the antimicrobial susceptibility testing, strain KP32558 and KP31166 were all CRKP and were resistant to all tested antibiotics (Table 1). Both strains exhibited the same MICs for tigecycline (16 mg/L) and colistin (256 mg/L). ECIM and mCIM tests indicated that the two strains produced metallo-β-lactamase.

**Table 1.** Antibiotic resistance characteristics (MICs, mg/L) of 4 clinical *K. pneumoniae* strains and the transconjugant of KP32558.

| Antibiotics | KP31166 | KP32558 | J53 | Transconjugant 1<br>(MCR-8.2) | Transconjugant 2<br>(NDM-5) |
| --- | --- | --- | --- | --- | --- |
| Amikacin | >=64 | >=64 | <=2 | <=2 | 4 |
| Tobramycin | >=16 | >=16 | <=1 | 4 | <=1 |
| Doxycycline | >=16 | >=16 | 2 | >=16 | 2 |
| Minocycline | >=16 | >=16 | 2 | 8 | 2 |
| Tigecycline<sup>a</sup> | 4 | 4 | 0.25 | 0.25 | 0.25 |
| Cefepime | >=32 | >=32 | <=0.12 | <=0.12 | 16 |
| Ceftazidime | >=64 | >=64 | <=0.12 | <=0.12 | >=64 |
| TZP | >=128 | >=128 | <=4 | >=128 | >=128 |
| SCF | >=64 | >=64 | <=8 | <=8 | >=64 |
| Aztreonam | >=64 | >=64 | <=1 | <=1 | <=1 |
| Imipenem | >=16 | >=16 | <=0.25 | <=0.25 | >=16 |
| Meropenem | >=16 | >=16 | <=0.25 | <=0.25 | >=16 |
| Levofloxacin | >=8 | >=8 | <=0.12 | 1 | <=0.12 |
| Ciprofloxacin | >=4 | >=4 | <=0.25 | 0.5 | <=0.25 |
| Colistin<sup>a</sup> | 256 | 256 | 1 | 4 | 1 |
| SXT | >=320 | >=320 | <=20 | >=320 | <=20 |

<sup>a</sup>MIC of tigecycline and colistin was detected using microdilution broth method. SAM, Ampicillin/sulbactam; TZP, Piperacillin/tazobactam; SCF, Cefoperazone/sulbactam; SXT, Sulfamethoxazole/trimethoprim.

Strain KP32558 was randomly selected for further analysis, which was not hypermucoviscous and was assigned to ST656. Neither *rmpA/rmpA2* genes nor *iucABCDiutA* and *iroBCDN* virulence gene clusters were found in this strain. Resistance gene annotation showed that more than 40 resistance genes were located on the chromosome and plasmids of *K. pneumoniae* isolate KP32558 (Table S1).

### Plasmid containing *bla*_NDM-5_ gene

*K. pneumoniae* KP32558 contained 8 plasmids with 310633 bp, 145073 bp, 109949 bp, 95351 bp, 46161 bp, 5721 bp, 4574 bp and 2927 bp in length. The GenBank accession numbers of these plasmids were CP076031-CP076038. The *bla*_NDM-5_ gene was found in the fifth plasmid pKP32558-5-ndm5 (46161 bp in size), which belonged to the IncX3 group and did not harbor any resistance genes other than *bla*_NDM-5_. Microbial nucleotide search for GenBank showed that pKP32558-5-ndm5 was nearly identical to a series of *bla*_NDM-5_-carrying plasmids, and they had the same size in length. Specifically, pKP32558-2-ndm5 showed 100% identity and coverage with plasmid pCREC-591_4 (CP024825, 46161 bp) from *E. coli* strain CREC-591, and 100% identity and 99.8% coverage with the plasmid pNDM_MGR194 (KF220657, 46253 bp), which was a typical *bla*_NDM-5_-carrying plasmid recovered from a *K. pneumoniae* isolate in India (25). Moreover, we found a genomic island region in pKP32558-5-ndm5, the *bla*_NDM-5_ gene and its flanking contents were also located in this region.

Similar to other *bla*_NDM-5_-carrying plasmid, several conjugal transfer genes (such as virD4, virB4, virB8 and virB9) were identified in plasmid pKP32558-5-ndm5. Conjugation experiment showed that this plasmid could be successfully transferred to the recipient *E. coli* J53 as previous report (26) at a frequency of 10^-4^ (transconjugant/recipient), and the transconjugant displayed resistance to imipenem and meropenem (Table 1).

### Presence of *mcr-8* gene in a two-replicons plasmid

An *mcr-8* gene (1698 bp) was found in the second plasmid pKP32558-2-mcr8 by blasting against the resfinder database, which was finally confirmed to be *mcr-8.2*. Microbial nucleotide search for GenBank suggested that pKP32558-2-mcr8 had the highest alignment score with the plasmid pKPNH54.1, which was derived from *K. pneumoniae* strain NH54, and did not contain *mcr* gene. Comparative genomic analysis was conducted among pKP32558-2-mcr8, pKPNH54 and other 10 *mcr-8.2*-containing plasmids obtained from NCBI database. The result showed that these plasmids were not identical to each other (Figure 2A), and the plasmids pKPNH54.1 (CP024917), pMCR8_020135 (CP037964) and pVNCKp115 (LC549807) could be aligned to different parts of pKP32558-2-mcr8. Despite the differences between these *mcr-8.2*-carrying plasmids, the commonality between them was that they had identical genetic environments around *mcr-8.2* gene, IS*Kpn26*-orf-*mcr-8.2*-IS*Ecl1-copR-baeS-dgkA* (Figure 2B). Besides, plasmid pZZW20-88k, pD120-1_83kb and p201936D-mcr8-345kb had a short fragment deletion in front of IS*903B* compared to other plasmids. Of note, these genes were located on a genomic island region.

**Figure 2.**
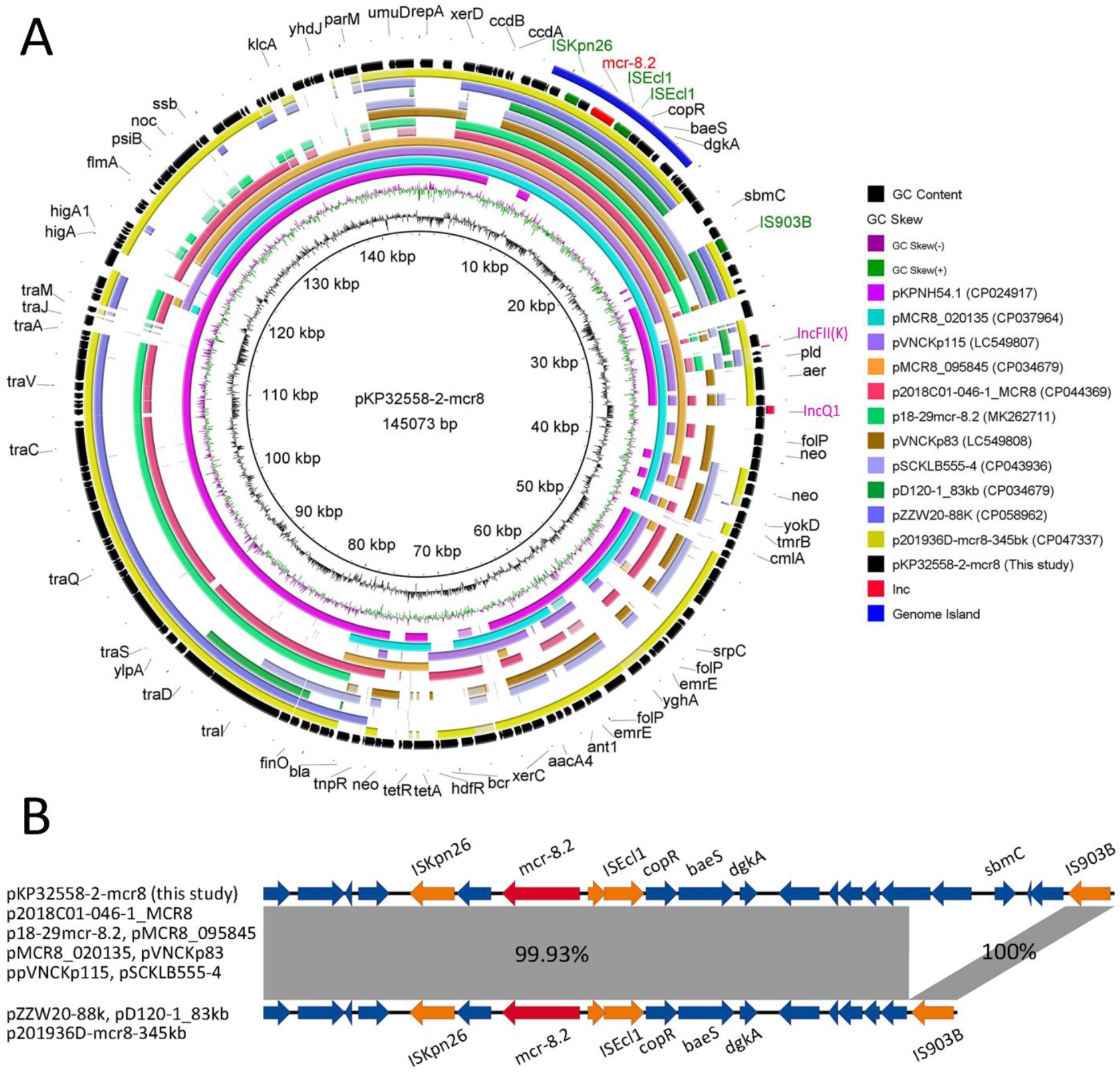
Genetic features of the mcr-8.2-carrying plasmid in *K. pneumoniae* strain KP32558. (A) Circular map of plasmid pKP32558-2-mcr8 and its reference plasmids. The outer blue circle is the genomic island region of plasmid pKP32558-2-mcr8. Red text on the plasmid map indicates *mcr-8.2* gene, green text indicates insertion sequences around *mcr-8.2*, and pink text represents the replicons. (B) Genetic contents of *mcr-8.2* gene. Coding sequences are indicated by arrows. Sequences of shared homology between two plasmids are marked by gray shading.

Plasmid typing suggested that these *mcr-8.2*-carrying plasmids contained 7 different replicons, and IncFII(K) replicon was found in most plasmids (9/11) (Figure 3). In addition to IncFII(K) replicon, the plasmid pKP32558-2-mcr8 also had a truncated IncQ1 type replicon, which was also present in pMCR8_020135 and pMCR8_095845. The two plasmids were derived from different hospitals of China. Phylogenetic tree showed that these plasmids were clustered into 3 major clades. Plasmid pKP32558-2-mcr8, pMCR8_020135, pMCR8_095845 and pVNCKp115 belonged to the same clade.

**Figure 3.**
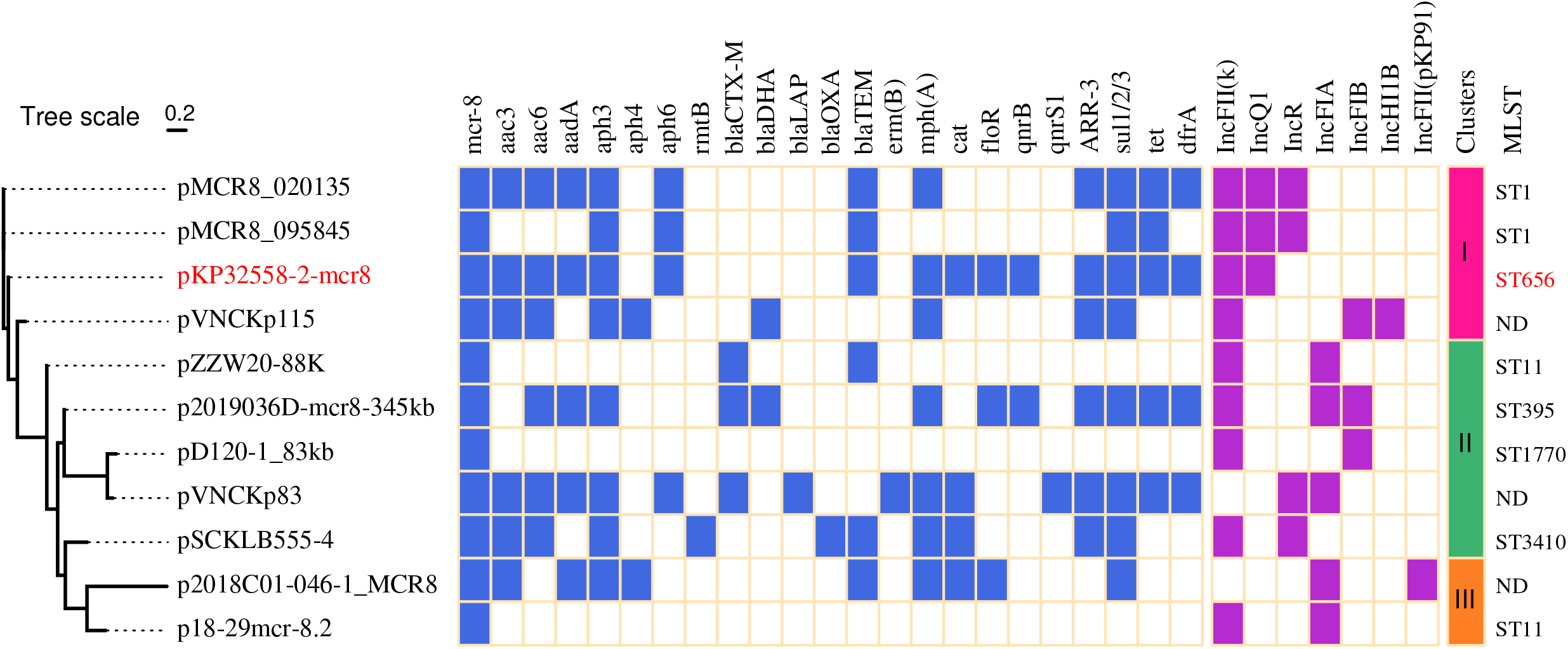
Resistance genes and replicons of mcr-8.2-carrying plasmids. The pKP32558-2-mcr8 (CP CP076032) was derived from *K. pneumoniae* KP32558 of this study. The genomic sequences of plasmids pMCR8_020135 (CP037964), pMCR8_095845 (CP031883), pVNCKp115 (LC549807), pZZW20-88K (CP058962), p2019036D-mcr8-345kb (CP047337), pD120-1_83kb (CP034679), pVNCKp83 (LC549808), pSCKLB555-4 (CP043936), p2018C01-046-1_MCR8 (CP044369) and p18-29mcr-8.2 (MK262711) were obtained from NCBI database.

### Resistance gene and conjugation experiment of plasmid pKP32558-2-mcr8

Resistance genes were identified using resfinder database. In addition to *mcr-8.2*, several other resistance genes were found in plasmid pKP32558-2-mcr8, including *aac, aad* and *aph* for aminoglycoside, *bla*_TEM_ for β-lactam, *mph(A)* for macrolide, *arr-3* for rifampicin, *sul1/2/3/* for sulphonamide, *tet* for tetracycline and *dfrA* for trimethoprim antibiotics (Figure 3). Conjugation experiments showed that the plasmid pKP32558-2-mcr8 could be transferred into *E. coli* J53 at a frequency of 10^-7^.

### MLST of *mcr*-8.2-carrying strains

MLST analysis suggested that these *mcr*-8.2-carrying strains belonged to different ST types, including ST1, ST656, ST11, ST395, ST1770 and ST3410. Among them, ST1, ST656 and ST3410 belonged to the same clonal complex, and there was only one allele difference between them. For three isolates, their chromosomal sequences were not deposited in GenBank, therefore the MLST results could not be obtained (Figure 3).

### Chromosomal factors account for colistin and tigecycline resistance

Two-component systems *PmrA*/*PmrB*, *PhoP*/*PhoQ* and *CrrA*/*CrrB*, and the regulator *mgrB* gene were analyzed to identify the potential mutations involved in colistin resistance. Genes in *K. pneumoniae* strain MGH 78578 (CP000647) were used as controls. The results showed that strain KP32558 contained amino acid substitution in *mgrB* (I45F), *pmrA* (D86E), *pmrB* (T246A and G345E) and *crrB* (C68S and V193G). PROVEAN was used for predicting the functional effect of amino acid substitution, indicating that 3 mutations in *mgrB, pmrA* and *crrB* (V193G) genes were deleterious (have a damaging effect on protein function), while the other 4 mutations were neutral (Table 2). These mutations might partly contribute to colistin resistance in *K. pneumoniae* strain KP32558.

**Table 2.**
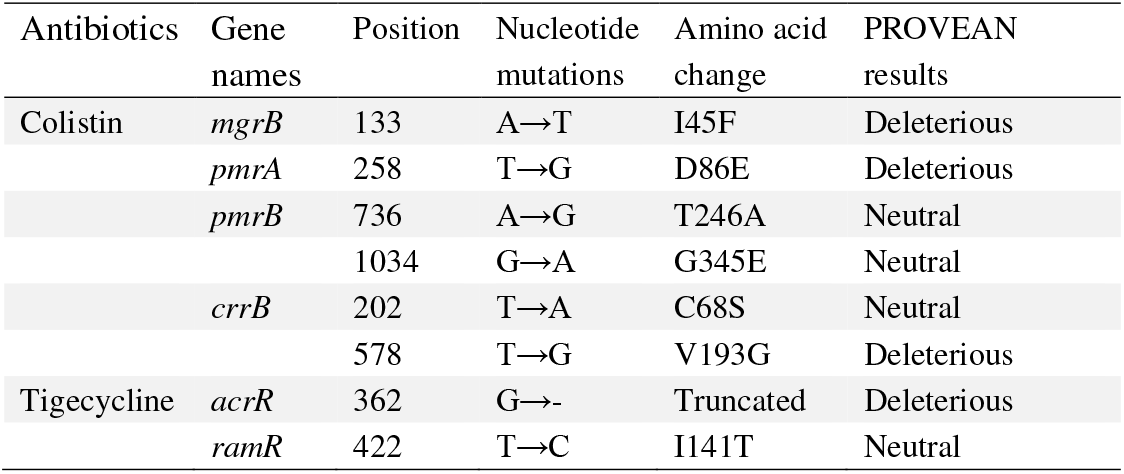
Mutations in chromosomal genes related to colistin and tigecycline resistance for *K. pneumoniae* KP32558

The efflux pump regulator genes *ramA, marR, soxR* and *acrR* of KP32558 were investigated for mutation using *K. pneumoniae* strain MGH 78578 as negative reference. A single-base deletion happened at position 362 of gene *acrR*, resulting in the premature termination of transcription, which could be associated with resistance to tigecycline. Another amino acid substitution was found in gene *ramR* (I141T), which had the neutral effect on the function.

### Growth curves of *K. pneumoniae* KP32558 and transconjugants

Compared to a carbapenem-resistant and hypervirulent *K. pneumoniae* isolate KP22937 (SAMN17245924) and a standard strain *K. pneumoniae* ATCC 700603, KP32558 had a slower growth rate. While the growth of transconjugant 1 (NDM-5) and transconjugant 2 (MCR-8.2) was almost indistinguishable from that of *E. coli* J53 (Figure 4).

**Figure 4.**
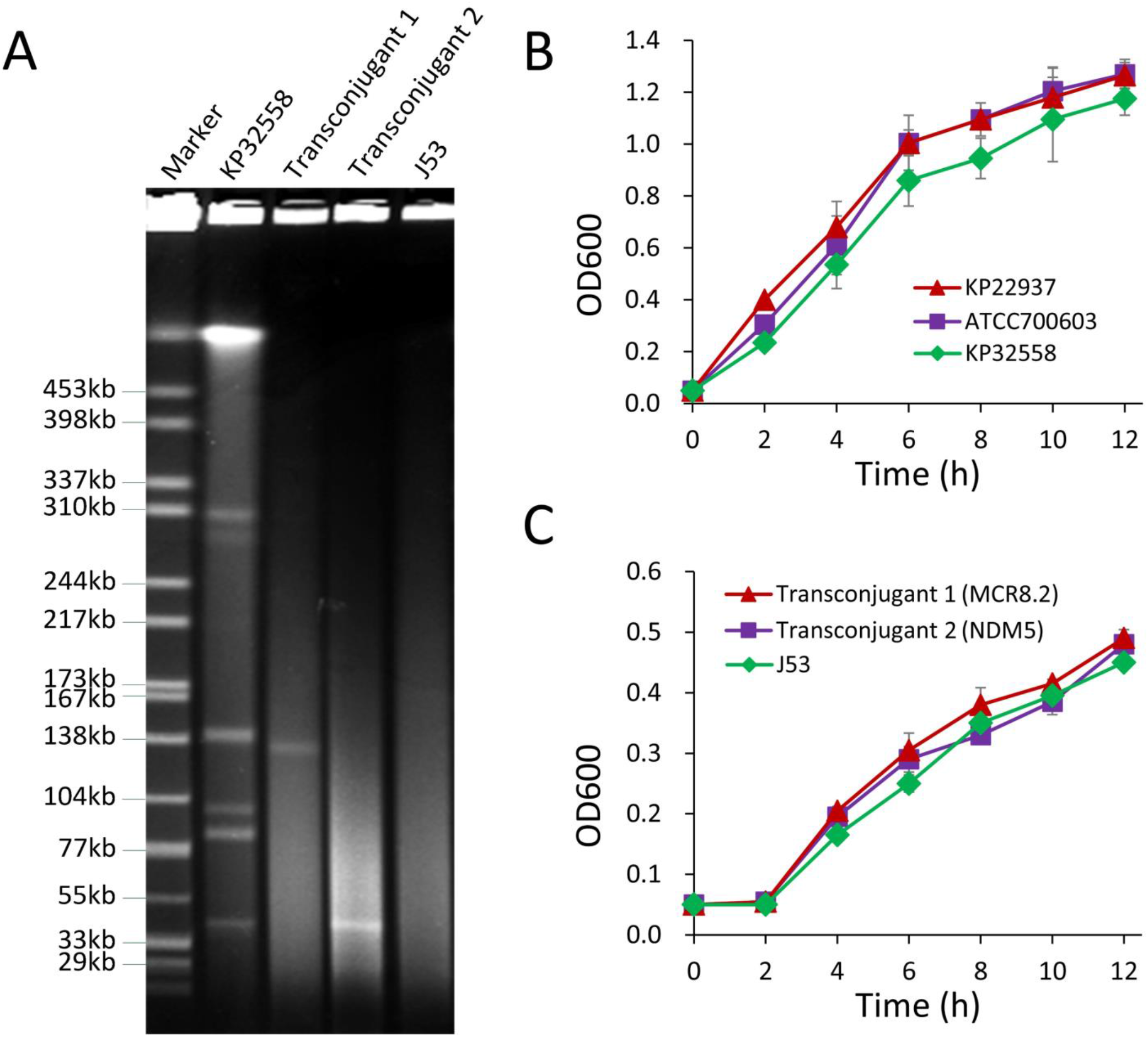
S1-PFGE profiles (A) and growth curves (B) of *K. pneumoniae* clinical strain, recipient bacterium *E. coli* J53 and transconjugants.

## DISCUSSION

The widespread of CRKP has aroused great attention worldwide. For CRKP that produces both NDM and extended-spectrum-β-lactamase (ESBL), neither ceftazidime/avibactam nor aztreonam can be used. Once these strains are resistant to colistin and tigecycline, treatment options are very limited. In this study, we identified an NDM-5-producing *K. pneumoniae* isolate recovered from a lung transplant patient, which simultaneously carried the plasmid-encoded *mcr-8.2* gene and chromosomal gene-mediated resistance to colistin and tigecycline. This represents the rare report of *K. pneumoniae* clinical isolate with multiple resistance mechanisms to the last-resort antimicrobials.

NDM had been spread in various species of bacteria in China (27). These genes are located on different types of plasmids, such as IncFII, IncIB and IncX3, among which IncX3 is the most reported. In our study, the NDM-5 gene was also located on IncX3 plasmid. IncX3 plasmids carrying NDM-5 gene are widely found in *Enterobacterales* bacteria, including *E. coli*, *K. pneumoniae, Proteus* spp. and *Klebsiella aerogenes* (28, 29), hinting its wide transmission.

Since the *mcr-8.1* gene from pig-derived *K. pneumoniae* strain KP91 was first reported in 2018, multiple *mcr-8* variants have been identified, ranging from *mcr-8.2* to *mcr-8.5* (30). Currently, *K. pneumoniae* producing both MCR-8 and NDM has emerged, such as *mcr-8.1* and *bla*_NDM-1_ (animal origin), *mcr-8.2* and *bla*_NDM-1_ (human origin), *mcr-8.1* and *bla*_NDM-5_ (human origin), and *mcr-8.5* and *bla*_NDM-5_ (animal origin) (30–33). In our study, we also identified the *K. pneumoniae* isolate co-harboring *mcr-8.2* and *bla*_NDM-5_ in lung transplantation ward, and should be alerted to its in-hospital spread.

Comparative genomic analysis was conducted to elucidate the genetic characteristics of *mcr-8.2*-carrying plasmid pKP32558-2-mcr8. This plasmid had two replicons, an intact IncFII(K) and a truncated IncQ1. Besides, eight of the ten *mcr-8.2*-carrying plasmids obtained from the GenBank database also had IncFII(K) type replicons, and the other two plasmids carried other IncF type replicons (Figure 3). This was consistent with the previous report that the IncF type plasmids are widely distributed in clinically relevant *Enterobacterales* isolates, and contribute to the fitness of bacterial host by providing antimicrobial resistance and virulence determinants (34). IncQ1 type plasmids are efficient antimicrobial resistance determinants among Gram-negative bacteria (35). The truncated IncQ1 replicon in the plasmid pKP32558-2-mcr8 was also observed in the other two plasmids, pMCR8_020135 and pMCR8_095845, indicating that these three plasmids may share a common ancestor. Phylogenic analysis of these *mcr-8.2*-carrying plasmids showed that plasmid pKP32558-2-mcr8, pMCR8_020135, pMCR8_095845 and pVNCKp115 were clustered into the same clade. At the same time, the latter three plasmids can match different parts of pKP32558-2-mcr8, indicating that there is a close genetic relationship between them.

MLST analysis showed that the *K. pneumoniae* strain KP32558 was assigned to ST656, a rare ST type and firstly reported in China in 2012 (36). The frequently detected carbapenemase in these bacteria was KPC-2 (37) and NDM-1 (38). Our study represents the first report of the emergence of NDM-5 and MCR in ST656 *K. pneumoniae* clinical isolate. The other 10 *mcr-8.2*-carrying plasmids came from bacteria with diverse ST types, including ST1, ST11, ST395, ST1770 and ST3410, suggesting the intra-species spread of mcr-8.2. It was noticeable that pMCR8_020135 and pMCR8_095845 were carried by ST1 *K. pneumoniae* strains, there was only one allele difference between ST656 and ST1, and they belonged to the same clonal complex. These results strengthened our hypothesis that the three plasmids share a common ancestor.

It is believed that without the antibiotic pressure, the acquisition of resistance genes in plasmids will impose fitness costs on their host (39). However, our recent study has shown that the growth of the transconjugant was almost indistinguishable from that of recipient *E. coli* EC600 (26), and another study also showed that up to 75.9% (22/29) *Enterobacterales* strains did not produce fitness costs after obtaining the IncX3 plasmid (40). The above results suggested that the acquisition of IncX3 plasmid might not confer a fitness cost to the host and then facilitate the rapid dissemination of the plasmid. In this study, the *bla*_NDM-5_-carrying IncX3 plasmid pKP32558-5-ndm5 could be transferred to *E. coli* J53 and the transconjugant had similar growth curve compared with *E. coli* J53, indicating that strain KP32558 probably acquired the resistance to carbapenems by obtaining the *bla*_NDM-5_-carrying and self-transmissible plasmid.

The MIC values for colistin in *mcr-8*-carrying isolates are usually between 4 and 32 mg/L (31, 41). While in this study, the MIC of *K. pneumoniae* strain KP32558 and transconjugant 2 were 256 and 16 mg/L, respectively, hinting that this strain had other colistin resistance mechanisms in addition to the *mcr-8.2* gene. Chromosomal mutations in genes encoding *PhoP/PhoQ, PmrA/PmrB* and *CrrA/CrrB* TCSs and mutations in *mgrB* gene can confer resistance to colistin in *K pneumoniae* (42). In isolate KP32558, a total of six amino acid substitutions were identified in *mgrB,pmrA,pmrB* and *crrB* genes, among which the mutations I45F in *mgrB*, D86E in *pmrA* and V193G in *crrB* were deleterious according to the PROVEAN tool. The D86E mutation has been described previously (43), while the other two mutations are reported for the first time. These results revealed that *mcr-8.2* gene and chromosomal TCS may have synergistic effects in mediating bacterial resistance to colistin, as reported in a previous study (44).

High-level expression of efflux pump is the main mechanism of bacterial resistance to tigecycline (45). Mutation in local transcriptional repressor *acrR* and global transcriptional activator *ramA* could result in the overexpression of AcrAB efflux pump (46). One previous study reported 26 tigecycline-nonsusceptible *K. pneumoniae* isolates, among which 23 (88.5%) had mutations in *ramR*, and 2 of the remaining 3 isolates contained a mutation in *acrR* and had a tigecycline MIC of 4 mg/L (47). In this study, a mutation in *ramR* (I141T) was found in *K. pneumoniae* strain KP32558, and this mutation had a neutral effect on its function, as consistent with a previous report (48). In addition, we also found a single-base deletion at position 362 of *acrR*, which resulted in the premature termination of transcription and showed the deleterious effect on the function. Therefore, we speculated that the mutation in *acrR* gene may contribute to the resistance of KP32558 to tigecycline.

Infections caused by multidrug-resistant *K. pneumoniae* can cause serious consequences for patients, especially those with compromised immune systems. We previously reported a *K. pneumoniae* isolate KP22937 with an NDM-5-producing plasmid and a hypervirulent plasmid, which caused the death of a lung transplant patient (26). Fortunately, in this study, although pan-drug resistant *K. pneumoniae* isolates were isolated twice in BALF, the bacteria did not cause serious infections in this lung transplant patient, and was eliminated without using specific antibiotics. Several mechanisms may contribute to the lower pathogenicity of KP32558. First, no classic hypervirulent genes were found in this strain, which limited its *in-vivo* dissemination. Second, the previous report showed that plasmid carriage often imposes the reduction in fitness to the host (49). In this study, KP32558 carried 8 plasmids, growth kinetics assay showed that KP32558 had a slower growth rate compared to the standard strain *K. pneumoniae* ATCC 700603 and the *K. pneumoniae* strain KP22937 we previously reported. Although the plasmids carrying NDM-5 or MCR-8.2 did not impose fitness costs on the recipient bacteria, the impact of the other 6 plasmids on the fitness of KP32558 could not be ruled out.

In conclusion, we report an MCR-8.2 and NDM-5-producing pan-drug resistant ST656 *K. pneumoniae* isolate recovered from the BALF specimen of a lung transplant patient. The *mcr-8.2* gene was located on a hybrid plasmid containing IncFII(K) and IncQ1 composition replicons, and which may play synergistic effects together with chromosomal mutations in mediating colistin resistance. Efflux pump repressor mutations were also found and may involve in the tigecycline resistance of *K. pneumoniae* isolate, suggesting that this bacterium has evolved and acquired a variety of complex resistance mechanisms to the last-resort antimicrobials. Therefore, close surveillance is urgently needed to monitor the prevalence of this clone.

## Acknowledgements

We would like to thank all the colleagues in the Laboratory of Clinical Microbiology and Infectious Diseases for their assistance.

